# Genomic characterization and functional insights of multidrug-resistant *Klebsiella pneumoniae* strain BCSIR-JUMIID

**DOI:** 10.1101/2024.11.27.625610

**Authors:** Mohammad Omar Faruk, Md. Murshed Hasan Sarkar, MD Ismail Hossain, Abdullah Al-Mamun Shabuj, Manotush Bhim, Kaniz Mehzabin, Sanjana Fatema Chowdhury, Showti Raheel Naser, Tabassum Mumtaz, Md Firoz Ahmed

## Abstract

*Klebsiella pneumoniae* is a prominent opportunistic pathogen associated with multidrug resistance (MDR) and high morbidity and mortality rates in healthcare settings. The emergence of strains resistant to last-resort antibiotics, such as colistin and carbapenems, poses significant therapeutic challenges. This study presents the complete genome analysis of the MDR strain *K. pneumoniae* BCSIR-JUMIID to elucidate its genetic architecture, resistance mechanisms, and virulence factors. The genome of *K. pneumoniae* BCSIR-JUMIID, isolated from a pharmaceutical wastewater in Dhaka, Bangladesh, was sequenced using next-generation sequencing technologies. Bioinformatics tools were employed for genome assembly, annotation, and functional analysis. Phylogenetic relationships were established through whole-genome comparisons. Antibiotic resistance genes, virulence factors, and mobile genetic elements were identified using the Comprehensive Antibiotic Resistance Database (CARD), ResFinder-4.5.0, Virulence Factors Database (VFDB), and various phage identification tools. The genome of *K. pneumoniae* BCSIR-JUMIID consists of 5,769,218 bp with a G+C content of 56.79%, assembled into 343 contigs. A total of 6,062 coding sequences (CDS), including 1,087 hypothetical proteins, 49 tRNA genes, and 4 rRNA genes, were identified. Key loci involved in capsular polysaccharide and O-antigen biosynthesis (KL150, KL107-D1, O3b) were detected. A diverse array of antibiotic resistance genes was uncovered, including those conferring resistance to beta-lactams, quinolones, and colistin. Phage analysis revealed the presence of multiple dsDNA bacteriophages, and CRISPR-Cas systems indicated robust phage defense mechanisms. The genomic analysis of *K. pneumoniae* BCSIR-JUMIID provides a detailed understanding of its resistance and virulence mechanisms, highlighting its potential for horizontal gene transfer and rapid adaptation. These findings underscore the necessity for continued surveillance and novel therapeutic strategies to combat MDR *K. pneumoniae* infections effectively.

## Introduction

*Klebsiella pneumoniae* is a Gram-negative, encapsulated bacterium that has emerged as a major nosocomial pathogen, causing a range of infections including pneumonia, bloodstream infections, and urinary tract infections (1,2).The increasing prevalence of multidrug-resistant (MDR) strains of *K. pneumoniae* has compounded the challenge of managing these infections, as these strains often display resistance to multiple antibiotic classes, including beta-lactams, aminoglycosides, quinolones, and even last-resort antibiotics such as colistin and carbapenems (2,3).

The global dissemination of MDR *K. pneumoniae* is primarily driven by its remarkable genetic plasticity, which allows the bacterium to acquire and disseminate antibiotic resistance genes through horizontal gene transfer mechanisms involving plasmids, transposons, and integrons (4).Furthermore, the presence of mobile genetic elements, such as bacteriophages and CRISPR-Cas systems, enhances the bacterium’s ability to evade antimicrobial treatments and adapt to various environmental pressures (18).

The growing threat posed by MDR *K. pneumoniae* underscores the need for a detailed understanding of its genetic and molecular basis of antibiotic resistance and virulence. Genomic studies offer valuable insights into these mechanisms, providing a foundation for the development of effective therapeutic and preventive strategies.

Pharmaceutical manufacturing facilities and their associated waste streams represent a unique ecological niche where antimicrobial resistance (AMR) genes can flourish (5).The complex interplay of selective pressures, including the use of antibiotics in production processes and the discharge of pharmaceutical effluents into the environment, fosters the evolution and dissemination of MDR bacteria (19). Despite this, studies characterizing MDR *K. pneumoniae* from pharmaceutical waste through whole-genome sequencing (WGS) remain limited.

In this study, we present a comprehensive genomic analysis of the MDR strain *K. pneumoniae* BCSIR-JUMIID, isolated from a clinical sample. By elucidating the genomic determinants of antimicrobial resistance and virulence, our study aims to provide insights into the genetic diversity, evolution, and dissemination of MDR *K. pneumoniae* in pharmaceutical environments. Furthermore, understanding the genetic mechanisms underpinning MDR in environmental isolates is crucial for devising effective strategies to mitigate the spread of resistance genes and safeguard public health (20).

Our study employs next-generation sequencing technologies and advanced bioinformatics tools to comprehensively analyze the genome of *Klebsiella pneumoniae* BCSIR-JUMIID. The primary objectives include assembling and annotating the genome to provide a detailed genetic blueprint of this MDR strain. We aim to identify and characterize antibiotic resistance genes, virulence factors, and mobile genetic elements, thereby elucidating the mechanisms underlying its resistance and pathogenicity. Additionally, we performed phylogenetic analyses to determine the evolutionary relationships and potential sources of horizontal gene transfer, offering insights into how these elements contribute to the bacterium’s adaptability and virulence. Through this comprehensive approach, we seek to better understand the genetic factors that enable *K. pneumoniae* BCSIR-JUMIID to thrive in various environments and develop resistance to multiple antibiotics. Through this comprehensive genomic analysis, we aim to shed light on the mechanisms underlying MDR in *K. pneumoniae* and contribute to the development of targeted therapeutic strategies and effective surveillance measures to combat the spread of this formidable pathogen.

### Methodology

#### Sample Collection and DNA Extraction

The Klebsiella pneumoniae strain BCSIR-JUMIID was isolated from pharmaceutical wastewater collected in Dhaka, Bangladesh, under standard microbiological laboratory conditions. Genomic DNA was extracted using the Qiagen DNeasy Blood & Tissue Kit (Qiagen, Germany), following the manufacturer’s protocol to obtain high-quality DNA suitable for sequencing applications.

#### Whole-Genome Sequencing

Whole-genome sequencing was performed on the Illumina MiniSeq platform (Illumina, San Diego, CA, USA). Paired-end sequencing libraries were prepared using the Nextera XT DNA Library Prep Kit (Illumina), targeting an average insert size of 150 bp. Raw sequencing reads were quality-checked using FastQC (v0.11.9) (22), and adapter sequences were trimmed using Trimmomatic (v0.39) (23).

#### Genome Assembly

The high-quality sequencing reads were assembled using the Genome Assembly Service provided by the Bacterial and Viral Bioinformatics Resource Center (BVBRC) and EDGE Bioinformatics tools. These platforms employ robust assembly algorithms to generate high-quality draft genomes. BVBRC offers integrated assembly pipelines that allow for contig generation, scaffolding, and quality evaluation (24). EDGE Bioinformatics facilitates a user-friendly genome assembly process, integrating tools such as SPAdes (25), and providing graphical interfaces for assembly quality control (26).

#### Taxonomic Phylogenetic Analysis

Multi-Locus Sequence Typing (MLST) was performed using the PubMLST database (27), confirming the taxonomic identity of the strain as Klebsiella pneumoniae based on allelic profiles of seven housekeeping genes. The Kaptive tool was used to identify capsular polysaccharide synthesis (cps) loci and O-antigen biosynthesis regions. The analysis identified loci KL150, KL107-D1, and O3b with high confidence (28). Phylogenetic analysis was conducted using the Type Strain Genome Server (TYGS) (29), assessing the evolutionary relationships between K. pneumoniae BCSIR-JUMIID and other reference strains, including ATCC 13883 and DSM 30104.

#### Pangenome Analysis

Fourteen reference genome sequences of the Klebsiella genus were retrieved from the NCBI Genome database. Pangenome analysis was performed using the Integrated Pathogen Genomics Analysis (IPGA) tool at the National Microbial Genomics Data Center (NMDC) was employed to evaluate the genetic diversity across various Klebsiella pneumoniae strains (30). The results confirmed a high degree of genomic similarity between the strain BCSIR-JUMIID and other reference K. pneumoniae strains.

#### Genome Annotation and Functional Insights

Genome annotation was performed using several platforms to ensure a comprehensive identification and functional characterization of the genes. The assembled genome of Klebsiella pneumoniae strain BCSIR-JUMIID was annotated using the Genome Annotation Service provided by BVBRC. This service facilitates automatic gene prediction, functional annotation, and identification of various genetic elements such as antimicrobial resistance genes, virulence factors, and mobile genetic elements.

Additionally, EDGE Bioinformatics offers a streamlined approach for genome assembly, annotation, and metabolic pathway identification. It integrates tools like Prokka (30) for rapid prokaryotic genome annotation and provides quality control metrics to ensure accuracy. For detailed genome visualization, Proksee (formerly CGView) was used, allowing comprehensive circular genome visualization and highlighting features such as coding sequences (CDS), RNA genes, and antimicrobial resistance loci (31). To further enrich the annotation process, we utilized the RAST (Rapid Annotation using Subsystems Technology) system at the National Microbial Pathogen Data Resource (NMPDR). RAST provides a robust annotation pipeline that assigns Enzyme Commission (EC) numbers, Gene Ontology (GO) terms, and KEGG pathway annotations to identified genes, offering insights into the functional roles of the genes in the genome (32). This tool was particularly useful for annotating metabolic pathways and other critical functional features within the K. pneumoniae genome.

#### Antibiotic Resistance and Virulence Factor Analysis

The Comprehensive Antibiotic Resistance Database (CARD) (33) and ResFinder-4.5.0 (34) were used to identify antibiotic resistance genes, categorized by efflux, target alteration, inactivation, and reduced permeability. BVBRC and EDGE Bioinformatics provided insights into virulence genes. The BacMet database was used to identify metal and biocide-resistant genes (35).

#### Mobile Genetic Elements and Bacteriophage Diversity

Bacteriophage sequences were identified using Phigaro (v2.3.0), PHASTER, and VirSorter2, classifying phage family members and analyzing prophage regions (36,37). Prediction of mobile genetic elements (MGEs) and horizontal gene transfer (HGT) events was conducted using Alien_Hunter (v1.7) (38) and mobileOG-db. Further analysis of MGEs was performed using MobileElementFinder and PlasmidFinder from the Center for Genomic Epidemiology (CGE) (39). For plasmid sequences in assembled scaffolds, geNomad from https://nmdc-edge.org/ was used (40).

#### Defense Mechanisms Against Phage Infections

Defense mechanisms against viral infections were identified using DefenseFinder (41), PADLOC (42), and CRISPRCasFinder (43). These tools were used to detect antiviral defense systems such as CRISPR-Cas and restriction-modification (R-M) systems, providing insights into the bacterial defense strategies against phage infections.

## Results

### Genome Assembly and Taxonomic Identification

The assembled genome of *Klebsiella pneumoniae* BCSIR-JUMIID was submitted to the Comprehensive Genome Analysis service at PATRIC. The assembly resulted in 343 contigs, with a total length of 5,769,218 bp and an average G+C content of 56.79% (Table 1). The genome assembly metrics indicate a good-quality genome, with a contig N50 of 32,077 and a contig L50 of 51.

**Table 1.** Assembly Details.

| Feature | Value |
| --- | --- |
| Contigs | 343 |
| GC Content | 56.79% |
| Contig L50 | 51 |
| Genome Length | 5,769,218 bp |
| Contig N50 | 32,077 |

**Table 2.** Annotated Genome Features.

| Feature | Count |
| --- | --- |
| CDS | 6,062 |
| tRNA | 49 |
| rRNA | 4 |
| Hypothetical proteins | 1,087 |
| Proteins with functional assignments | 4,975 |
| Proteins with EC number assignments | 1,515 |
| Proteins with GO assignments | 1,251 |
| Proteins with Pathway assignments | 1,097 |
| Proteins with PATRIC genus-specific family (PLfam) assignments | 5,771 |
| Proteins with PATRIC cross-genus family (PGfam) assignments | 5,862 |

EDGE bioinformatics tools, including gottcha-speDB-b, gottcha2-speDB-b, gottcha-speDB-v, kraken2, pangia, and centrifuge, were employed to classify the sequencing reads. Each tool uses different algorithms and databases to provide a comprehensive view of the dataset’s taxonomic composition. A heatmap visualization of the results revealed that Klebsiella pneumoniae is the most abundant species in the BCSIR-JUMIID sample, as indicated by the darker red cells (Fig 1a). This finding was corroborated by whole-genome sequencing and multi-locus sequence typing (MLST), which confirmed Klebsiella pneumoniae with 100% certainty. Fifty-four exact matches were found in the PubMLST database, confirming the species. Taxonomically, the strain belongs to the phylum Pseudomonadota, class Gammaproteobacteria, order Enterobacterales, family Enterobacteriaceae, genus Klebsiella, and species Klebsiella pneumoniae. This bacterium is a significant pathogen in clinical settings, commonly associated with infections such as pneumonia, bloodstream infections, and urinary tract infections (1,2). The Kaptive analysis of the bacterial assembly “BCSIR-JUMIID” identified three main loci: KL150, KL107-D1, and O3b. The KL150 locus showed high match confidence with 98.19% coverage and 99.51% identity, revealing 19 out of 21 expected genes, indicating a nearly complete capsular polysaccharide synthesis region (Fig 1b). In contrast, the KL107-D1 locus displayed lower confidence with 72.73% coverage and 76.74% identity, containing only 4 of the 13 expected genes, suggesting a partial or altered capsular locus. The O3b locus exhibited high match confidence with 99.43% coverage and 98.85% identity, with 7 of 8 expected genes present, indicating a well-preserved O-antigen biosynthesis region. These findings highlight the genetic diversity and potential virulence mechanisms of the bacterial strain, providing insights crucial for understanding its pathogenicity and informing treatment strategies.

**Fig 1.**
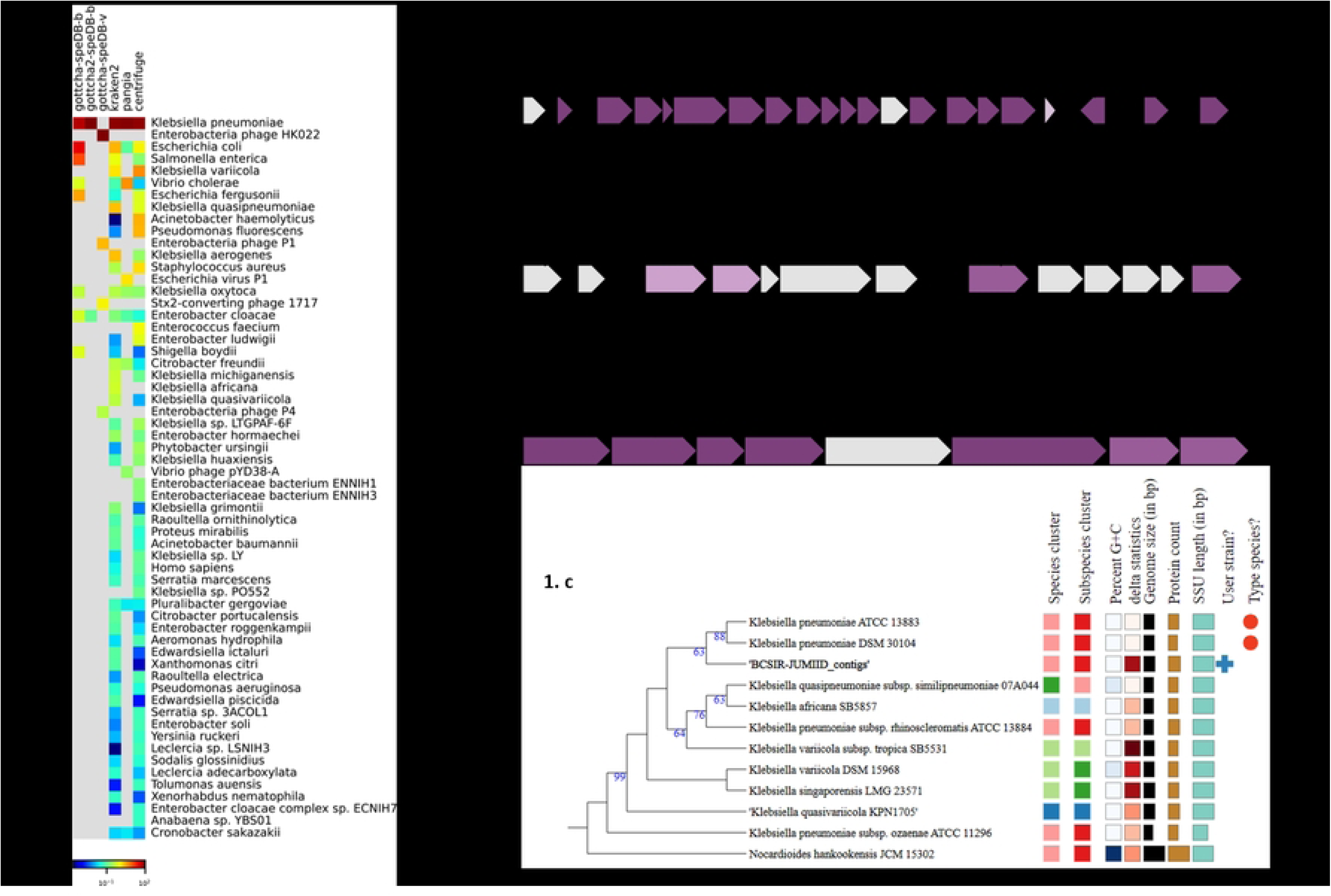
Taxonomic identification and genomic features of *Klebsiella pneumoniae* BCSIR-JUMIID. **(a)** Heatmap visualization of taxonomic composition based on the classification of sequencing reads using EDGE bioinformatics tools. The heatmap shows that *Klebsiella pneumoniae* is the most abundant species in the BCSIR-JUMIID sample. **(b)** Kaptive analysis showing the three main loci: KL150, KL107-D1, and O3b. **(c)** Phylogenetic tree generated using the TYGS platform reveals that *Klebsiella pneumoniae* BCSIR-JUMIID is closely related to *Klebsiella pneumoniae* strains ATCC 13883 and DSM 30104.

The evolutionary relationships among various *Klebsiella* species, including *Klebsiella* BCSIR-JUMIID, were elucidated using the TYGS platform (https://tygs.dsmz.de/). The phylogenetic tree generated reveals that *Klebsiella* BCSIR-JUMIID is a member of the *Klebsiella pneumoniae* family, a group of bacteria commonly associated with pneumonia (Fig 1c). Notably, *Klebsiella* BCSIR-JUMIID exhibits close genetic relatedness to *Klebsiella pneumoniae* strains ATCC 13883 and DSM 30104.

### Pangenome Analysis

The pangenome analysis of fifteen *Klebsiella* genomes, including *Klebsiella* BCSIR-JUMIID, revealed substantial genetic diversity and a highly dynamic pangenome structure. A total of 14,273 pan-gene clusters were identified, representing the cumulative genetic content of all genomes in the study. In contrast, the core genome—shared by all strains—consisted of only 569 core gene clusters, highlighting that while *Klebsiella* species share a conserved set of essential genes, the majority of the genome comprises accessory genes that contribute significantly to genetic variability and adaptation potential within the genus (Fig 2a). Functional classification of the core genome using COG (Clusters of Orthologous Groups) annotation revealed that out of the 569 core gene clusters, 1,171 genes were associated with metabolism, 626 with cellular processes and signaling, and 426 with information storage and processing. Additionally, 416 genes were either poorly characterized or unannotated. These core genes likely perform essential functions necessary for maintaining basic cellular processes, underlining their evolutionary conservation across Klebsiella species. (Fig 2b). Notably, the accessory genome of Klebsiella BCSIR-JUMIID contained several unique genes that are potentially associated with virulence. Genes encoding the preprotein translocase subunit SecA, signal peptidases, and ABC transport systems were identified.

**Fig 2.**
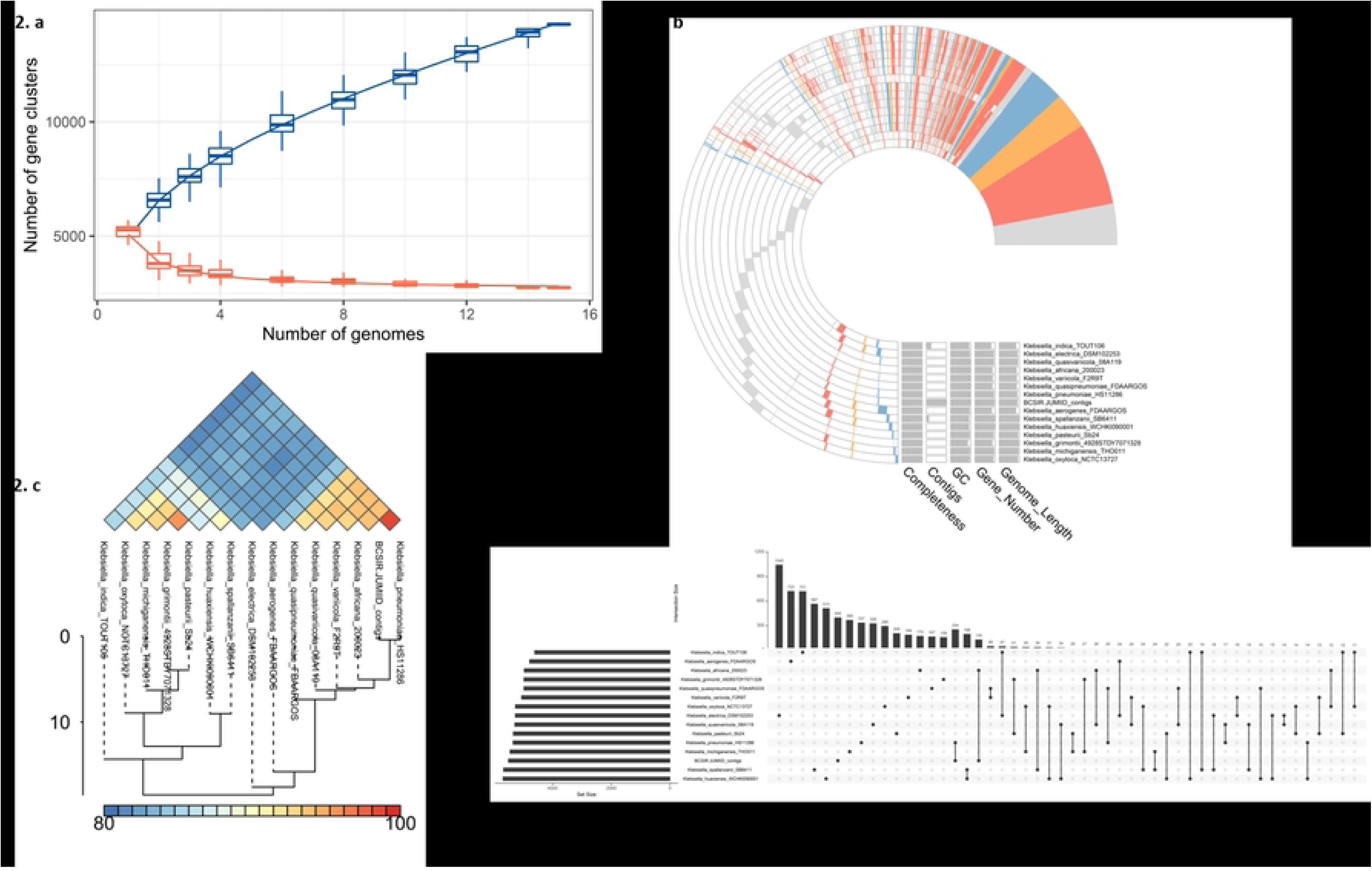
Pangenome analysis and genetic relationships of *Klebsiella* genomes, including *Klebsiella pneumoniae* BCSIR-JUMIID. **(a)** Pangenome and core genome distribution of *Klebsiella* species. **(b)** Functional classification of the core genome based on COG (Clusters of Orthologous Groups) annotation. **(c)** Average Nucleotide Identity (ANI) analysis of *Klebsiella pneumoniae* BCSIR-JUMIID and related strains. BCSIR-JUMIID exhibited the highest ANI similarity (99.60%) with *Klebsiella pneumoniae* HS11286, confirming its close evolutionary relationship within the *K. pneumoniae* species complex **(d)** UpSet plot illustrating the genetic diversity of *Klebsiella* strains.

The Average Nucleotide Identity (ANI) analysis provided additional insights into the genetic relationships among the strains. *Klebsiella* BCSIR-JUMIID exhibited the highest ANI similarity (99.60%) with *Klebsiella pneumoniae* HS11286, confirming its close evolutionary relationship within the *K. pneumoniae* species complex. ANI values above 95% typically indicate that genomes belong to the same species, supporting the classification of BCSIR-JUMIID as a strain of *K. pneumoniae*. BCSIR-JUMIID also showed high ANI values with *Klebsiella variicola* F2R9T (94.32%) and Klebsiella africana 200023 (94.96%), indicating close evolutionary relationships, though more distant than with K. pneumonia (Fig 2c). In contrast, ANI values with other species, such as Klebsiella aerogenes (84.26%) and Klebsiella grimontii (83.07%), were substantially lower, reflecting greater genetic divergence. An upset plot further illustrated the extent of this genetic diversity, showing that the number of unique gene clusters per genome ranged from 159 to 1,045, underscoring the variability in the accessory genomes of these strains (Fig 2d).

### Genome Annotation and Functional Insights

The genome of Klebsiella pneumoniae BCSIR-JUMIID was annotated using both EDGE Bioinformatics and RASTtk, yielding a comprehensive view of its functional and structural features. RASTtk annotated 6,062 CDS, comprising 1,087 hypothetical proteins and 4,975 functionally characterized proteins, suggesting substantial consistency between the two platforms regarding core functional annotations, though with variation in the number of hypothetical proteins (Fig 3a. The RASTtk analysis annotated genome features include 6,062 protein-coding sequences (CDS), 49 transfer RNA (tRNA) genes, and 4 ribosomal RNA (rRNA) genes, with no partial CDS, miscellaneous RNA, or repeat regions (Table 3a).

**Fig 3.**
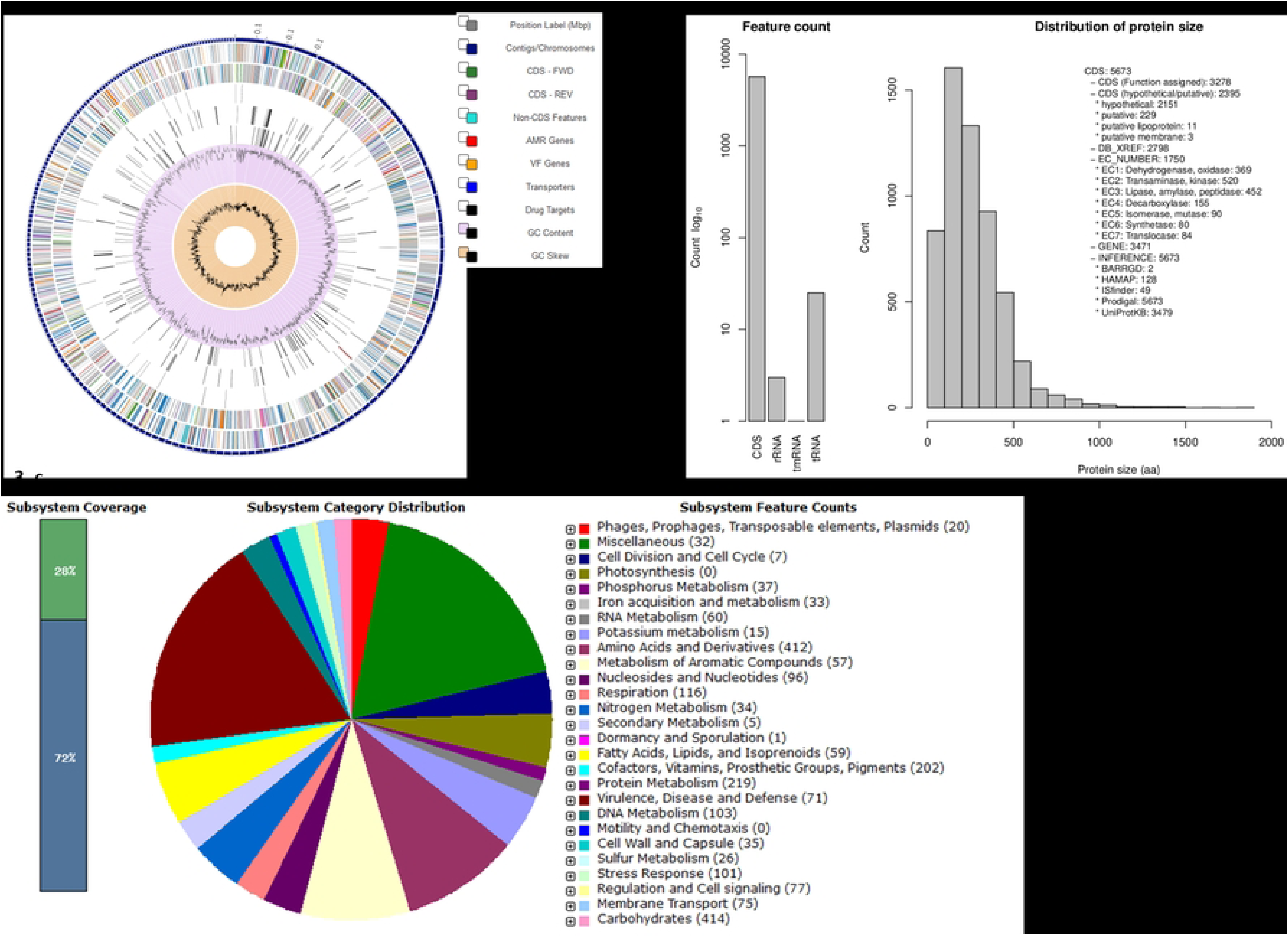
Genomic annotation and functional analysis of *Klebsiella pneumoniae* BCSIR-JUMIID using RASTtk and EDGE Bioinformatics. **(a)** A circular graphical display of the distribution of the genome annotations by Patric. This includes, from outer to inner rings, the contigs, CDS on the forward strand, CDS on the reverse strand, RNA genes, CDS with homology to known antimicrobial resistance genes, CDS with homology to know virulence factors, GC content and GC skew. The colors of the CDS on the forward and reverse strand indicate the subsystem that these genes belong to (see Subsystems below). **(b)** Insertion sequence and protein family classification identified by EDGE Bioinformatics. A total of 49 insertion sequences and extensive protein family classifications demonstrate the isolate’s genomic diversity and adaptability. **(c)** Subsystem analysis of *Klebsiella pneumoniae* BCSIR-JUMIID using RASTtk.

Conversely, EDGE Bioinformatics identified 5,673 coding sequences (CDS), including 2,395 hypothetical proteins, and functionally assigned 3,278 CDS. In addition, EDGE Bioinformatics identified 49 insertion sequences, indicating a high potential for horizontal gene transfer, a critical factor in genomic adaptability and the acquisition of antimicrobial resistance genes (Fig 3b). EDGE further classified 5,771 proteins into genus-specific protein families (PLfams) and 5,862 into cross-genus protein families (PGfams), demonstrating significant genomic diversity and adaptability. A detailed subsystem analysis of the genome using RASTtk revealed dominant functional categories, including carbohydrate metabolism, amino acid metabolism, protein metabolism, and virulence, disease, and defense mechanisms. The large proportion of genes in these subsystems indicates a broad metabolic capacity, allowing the bacterium to utilize a wide range of energy sources and thrive in diverse environments. Carbohydrate metabolism was one of the largest subsystems, underscoring the bacterium’s ability to metabolize sugars and polysaccharides, a key factor in its ecological flexibility (Fig 3c).

Metabolic pathways related to nitrogen, phosphorus, and sulfur metabolism were also identified, further emphasizing the bacterium’s adaptability to nutrient-limited environments, which may include clinical or environmental niches with fluctuating nutrient availability. Additionally, RASTtk analysis revealed the presence of regulatory mechanisms through genes involved in cell signaling and regulation, suggesting that Klebsiella pneumoniae BCSIR-JUMIID can fine-tune its gene expression in response to environmental stimuli, which may contribute to its survival in diverse and potentially challenging environments.

### Pathogenicity, Antibiotic Resistance and Virulence Island

The analysis of Klebsiella BCSIR-JUMIID isolates using the IslandPath-DIMOB tool has unveiled a diverse collection of proteins, enzymes, transporters, and regulatory elements, many of which are associated with resistance mechanisms, mobile genetic elements, and other essential bacterial processes. Utilizing tools like SIGI-HMM, which employs Hidden Markov Models to detect signatures of genomic islands, phages, and resistance genes, this analysis highlights critical genomic features (Figure 4a).

**Fig 4.**
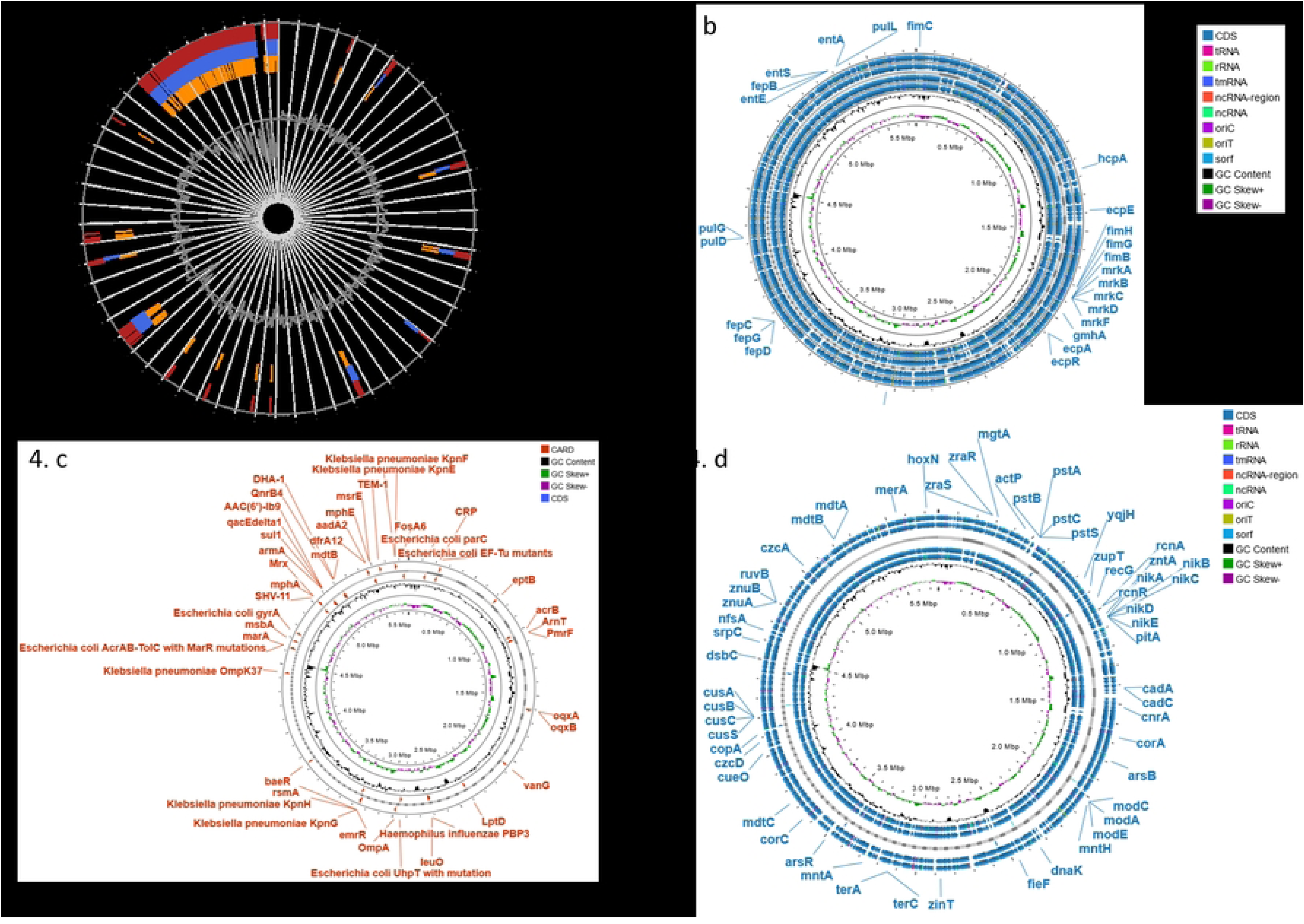
Comprehensive genomic analysis of *Klebsiella pneumoniae* BCSIR-JUMIID isolate, highlighting mobile genetic elements, virulence factors, antibiotic resistance mechanisms, and heavy metal resistance genes. **(a)** Genomic island and resistance gene identification IslandViewer 4. **(b)** Virulence factor analysis of *K. pneumoniae* BCSIR-JUMIID, identifying genes associated with pilus production, iron acquisition, biofilm formation, lipopolysaccharide (LPS) biosynthesis, and type II secretion system (T2SS). **(c)** Antibiotic resistance gene profile obtained CARD databases, illustrating resistance to aminoglycosides, fluoroquinolones, beta-lactams, macrolides, sulfonamides, tetracyclines, and polymyxins. **(d)** Heavy metal resistance gene analysis, identifying genes involved in arsenic, cobalt, zinc, cadmium, copper, and mercury detoxification and efflux, indicating the isolate’s capacity to survive in metal-contaminated environments.

The genomic analysis of Klebsiella pneumoniae BCSIR-JUMIID reveals an array of virulence factors that significantly enhance its pathogenic potential. The presence of a substantial set of genes including ecpA, ecpB, ecpC, ecpD, ecpE, and ecpR—underscores the isolate’s capacity for pilus production, essential for adhesion to host cells and biofilm formation. Iron acquisition is another critical aspect of the virulence profile of K. pneumoniae BCSIR-JUMIID. The isolate possesses genes such as entA, entB, entE, and entS are involved in enterobactin production, while fepB, fepC, fepD, and fepG facilitate its uptake. The fim operon, including the fim and mrk operons, encodes components of type 1 fimbriae, which facilitate attachment to host epithelial cells, particularly in the urinary tract, while the mrk operon encodes type 3 fimbriae essential for biofilm development on various surfaces. The biosynthesis of lipopolysaccharide (LPS) is another important virulence factor, with genes such as gmhA/lpcA, kdsA, and lpxC contributing to its production. Moreover, the ppdD/hcpA gene encodes fimbrial subunits that further enhance the bacterium’s adhesion to host tissues. The presence of genes from the Pul family, including pulD, pulG, pulI, pulK, and pulO, indicates the existence of a type II secretion system (T2SS), which secretes enzymes capable of degrading host polysaccharides (Figure 4b).

The whole genome analysis of *Klebsiella pneumoniae* strain BCSIR-JUIIMD, conducted using the Comprehensive Antibiotic Resistance Database (CARD) and ResFinder-4.5.0 tools, identified a broad array of antibiotic resistance genes and mechanisms. The ResFinder-4.5.0 analysis revealed resistance to multiple aminoglycosides mediated by genes such as armA, aac(6’)-Ib3, and aadA2. The strain also showed resistance to spectinomycin (aadA2), ciprofloxacin, and nalidixic acid (OqxB, OqxA, qnrB4), and various beta-lactams. Additionally, resistance was observed to sulfamethoxazole (sul1), trimethoprim (OqxB, OqxA, dfrA12), fosfomycin (fosA6), macrolides conferred by mph(A), msr(E), mph(E), tetracyclines mediated by tmexD3, TOprJ3, tmexC3, and streptogramins through msr(E). Chloramphenicol resistance was also detected (OqxB, OqxA). The CARD analysis identified additional resistance mechanisms, including antibiotic efflux pumps and target alteration. The eptB and ArnT genes were linked to resistance to peptide antibiotics, particularly polymyxin B and colistin, through target alteration. The Shigella flexneri acrA gene indicated resistance to multiple classes, including fluoroquinolones, cephalosporins, glycylcyclines, penams, tetracyclines, rifamycins, and phenicols, via an RND efflux pump. Perfect hits for oqxA and oqxB confirmed resistance to fluoroquinolones, glycylcyclines, tetracyclines, diaminopyrimidines, and nitrofuran antibiotics through an RND efflux mechanism. The LptD gene, conferred resistance to carbapenems, peptide antibiotics, aminocoumarins, and rifamycins via an ATP-binding cassette (ABC) efflux pump. The SHV-11 gene was identified as a beta-lactamase conferring resistance to carbapenems, cephalosporins, and penams. Other significant resistance genes included mphA and mphE, armA, and sul1. Reduced permeability mechanisms were indicated by the OmpA and *Klebsiella pneumoniae* OmpK37genes, linked to resistance to peptide antibiotics and beta-lactams, respectively. Additionally, mutations in Escherichia coli parC (S80I mutation) and gyrA (S83I mutation) genes conferred resistance to fluoroquinolones through target alteration (Fig 4c).

The analysis revealed several heavy metal resistance genes, indicating the isolate’s ability to thrive in contaminated environments. Genes such as ArsA, ArsB, and ArsH are involved in arsenic detoxification, while CzcD provides resistance to cobalt, zinc, and cadmium. Furthermore, copper resistance is conferred by the genes CusA, CusB, CusR, and PcoE, which facilitate the detoxification and efflux of copper ions. Additionally, MerD, MerE, and MerT are implicated in mercury detoxification, demonstrating the isolate’s potential for environmental survival under toxic metal stress. Additionally, the presence of copper and silver efflux systems, including CusC, CusA, and CusR, further reinforces the isolate’s ability to manage metal toxicity (Fig 4d).

### Bacteriophage Diversity, Mobile Genetic Elements and Horizontal Gene Transfer

The analysis of the Klebsiella pneumoniae isolate using mobileOG-db classified 637 protein families associated with mobile genetic elements (MGEs) into five major functional categories. These categories included Integration/Excision (IE) with 138 protein families, Replication/Recombination/Repair (RRR) with 174 protein families, Phage (P) with 124 protein families, Stability/Transfer/Defense (STD) with 81 protein families, and Transfer (T) with 120 protein families. The presence of these diverse protein families suggests that the isolate harbors a broad range of MGEs, such as conjugative plasmids, transposons, and bacteriophages, which likely contribute to its multidrug resistance (MDR) and virulence. Further investigation of the specific protein families identified through mobileOG-db is necessary to better understand the mechanisms driving these characteristics (Fig 5a).

**Fig 5.**
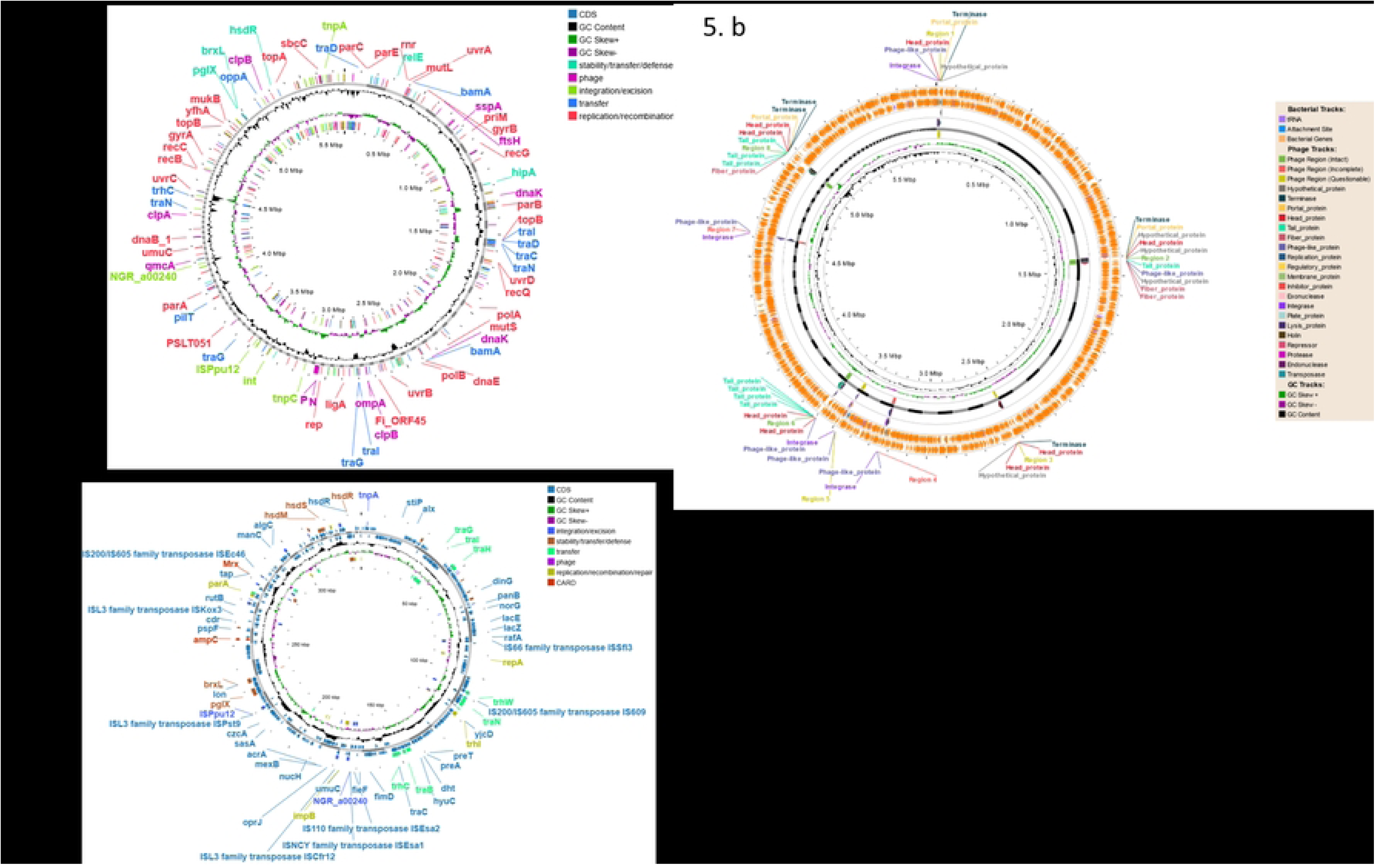
Characterization of mobile genetic elements (MGEs) and associated resistance and virulence factors in *Klebsiella pneumoniae* BCSIR-JUMIID. **(a)** mobileOG-db classification of 637 protein families linked to MGEs, categorized into five functional groups: Integration/Excision (IE), Replication/Recombination/Repair (RRR), Phage (P), Stability/Transfer/Defense (STD), and Transfer (T). **(b)** Bacteriophage analysis using PHASTER revealing a range of prophage regions, including intact, questionable, and incomplete prophages. **(c)** geNomad analyses showing multiple plasmid replicons harboring antibiotic resistance genes and virulence factors.

A detailed bacteriophage analysis using Phigaro 2.3.0, PHASTER, and VirSorter2 revealed a diverse array of bacteriophages with varied structural and functional properties. The analysis identified phages from the families Siphoviridae, Myoviridae, and Podoviridae, suggesting a combination of lysogenic (Siphoviridae) and lytic (Myoviridae) cycles, as well as diverse DNA injection mechanisms (Podoviridae). PHASTER analysis identified six prophage regions, including a questionable 19.9 Kb region linked to PHAGE_Shigel_SfIV, and intact regions of 36 Kb, 27.1 Kb, and 24.1 Kb associated with PHAGE_Edward_GF_2, PHAGE_Klebsi_ST512_KPC3phi13.2, and PHAGE_Pseudo_phiPSA1, respectively. Additionally, two incomplete prophage regions (18.6 Kb and 11.2 Kb) were linked to PHAGE_Escher_phiV10 and PHAGE_Escher_HK639. These findings suggest that the isolate may possess a significant number of prophages that could play a role in the horizontal transfer of resistance and virulence genes, thus enhancing the pathogenic potential of the strain (Fig 5b).

The MobileElementFinder and PlasmidFinder tools were used to detect plasmid replicons within the isolate. Several plasmid types were identified, including IncC, IncFIB(pNDM-Mar), IncHI1B(pNDM-MAR), IncR, IncY, and repB(R1701). These plasmids are well known for their association with the dissemination of antibiotic resistance genes. The use of the geNomad workflow for scaffold analysis revealed multiple plasmids carrying important resistance genes, including qnrB4, blaDHA-1, blaTEM-1B, blaSHV-182, OqxA, OqxB, fosA, mph(A), qacE, aac(6’)-Ib-cr, mph(E), msr(E), dfrA12, and aadA2. These genes confer resistance to a broad range of antibiotics, including quinolones, beta-lactams, and aminoglycosides. Additionally, virulence factors such as iutA, mrkA, terC, and fimH were also identified, indicating the potential for enhanced pathogenicity (Fig 5c).

### Defense Mechanisms against Phage Infections

The bacterial defense systems of *Klebsiella* BCSIR-JUMIID were comprehensively characterized using three complementary bioinformatics tools: PADLOC, DefenseFinder, and CRISPRCasFinder 4.2.20. These analyses revealed a diverse and multi-layered defense strategy involving CRISPR-Cas systems, restriction-modification (RM) systems, abortive infection (Abi) mechanisms, and less characterized defense elements, along with phage-encoded antidefense systems, which highlight the ongoing evolutionary dynamics between bacteria and bacteriophages.

CRISPR-Cas systems were detected consistently by both DefenseFinder (Fig 6a) and PADLOC (Fig 6b), which identified a Class 1, Subtype IV-A CRISPR-Cas system known for targeting phage DNA. CRISPR arrays were also confirmed by CRISPRCasFinder (Fig 6c), with a particularly strong prediction on Contig 41 (evidence level 3), suggesting a likely functional CRISPR-Cas defense. Additionally, Cas proteins necessary for viral DNA targeting were found across multiple contigs, including Type III-A on Contig 71 and Type U on Contig 41, indicating that *Klebsiella* BCSIR-JUMIID harbors several functional CRISPR-Cas systems.

**Fig 6.**
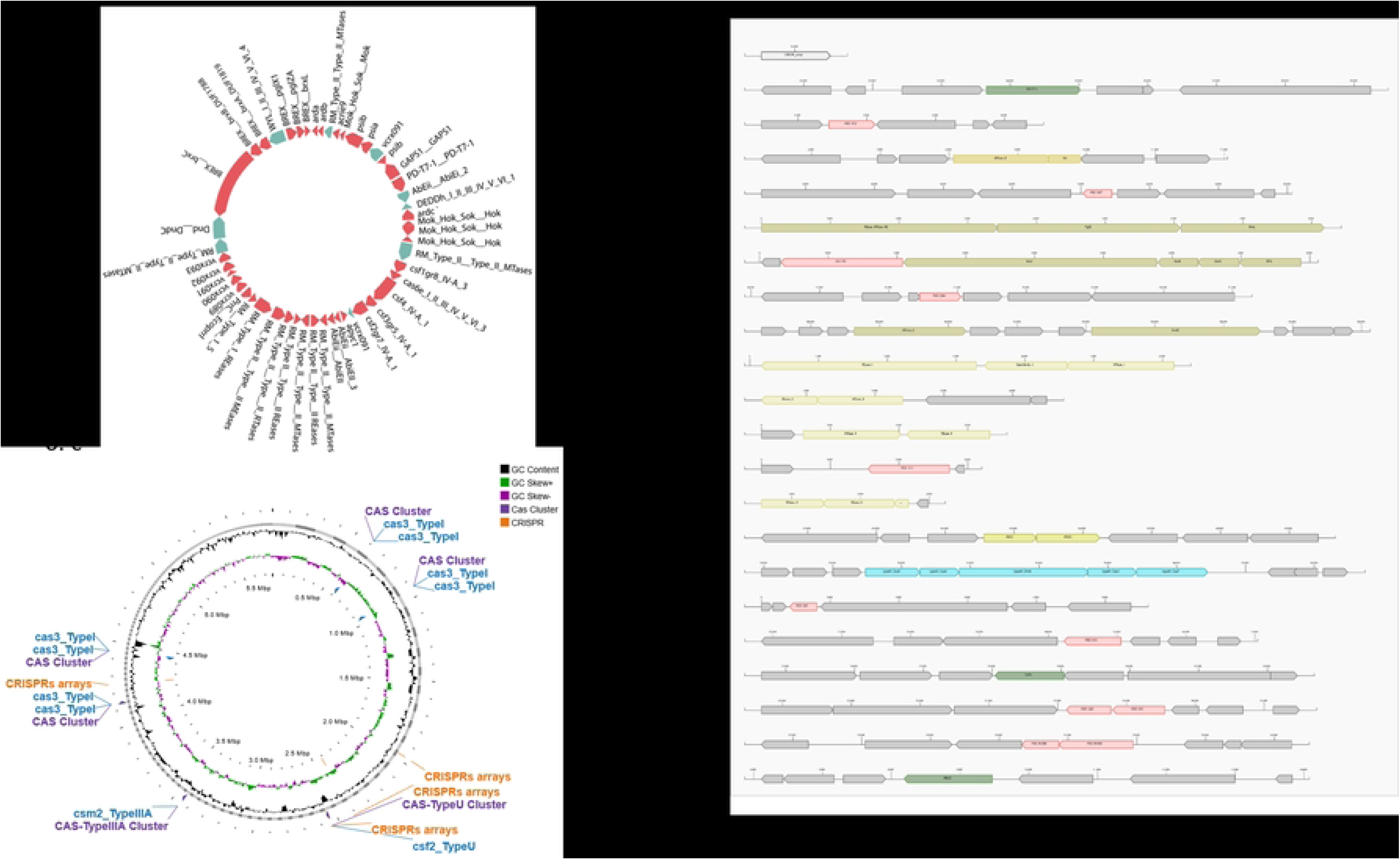
Distribution of defense systems across *Klebsiella pneumoniae* BCSIR-JUMIID. (a) DefenseFinder results highlighting key bacterial defense systems, CRISPR-Cas, restriction-modification (RM) systems, abortive infection systems, BREX, GAPS1, and toxin-antitoxin systems. (b) PADLOC results showing a comprehensive view of the bacterial defense systems identified, including CRISPR arrays, RM systems, the BREX system, and other systems. (c) CRISPRCasFinder results detailing the CRISPR-Cas system analysis. The tool identified multiple CRISPR arrays and Cas protein clusters.

Both PADLOC and DefenseFinder identified multiple restriction-modification (RM) systems, including Type I and Type II systems, which play a critical role in protecting bacterial genomes by cleaving foreign DNA and methylating specific sequences. These systems are key elements in *Klebsiella* BCSIR-JUMIID’s arsenal against bacteriophages, enhancing its capacity to resist phage infection.

In addition to CRISPR-Cas and RM systems, the BREX (I) system was detected by both PADLOC and DefenseFinder, suggesting that *Klebsiella* BCSIR-JUMIID employs this non-cleaving phage defense mechanism to inhibit viral replication. The detection of abortive infection (Abi) systems, including AbiE and AbiU, further adds to the multilayered nature of the bacterial defense, providing an additional fail-safe mechanism by inducing programmed cell death in infected cells to prevent the spread of phages.

Furthermore, both PADLOC and DefenseFinder identified several other less-studied or novel defense systems. PADLOC detected PD-T7-1, PDC-S12, and VSPR systems, along with BrxL and BrxC proteins associated with the BREX system, whereas DefenseFinder highlighted the GAPS1 system and the Mok-Hok-Sok toxin-antitoxin system, which may contribute to bacterial stress responses and phage resistance. Of particular interest was the identification by DefenseFinder (Fig 6a) of phage-encoded antidefense systems, including Anti-RM systems (e.g., Ard proteins) and Anti-CRISPR systems (e.g., acrIE9), which allow phages to evade bacterial immune defenses. This highlights the ongoing evolutionary arms race between *Klebsiella* BCSIR-JUMIID and bacteriophages, as the phages continue to evolve mechanisms to overcome bacterial defense strategies.

## Discussion

Multidrug-resistant (MDR) *Klebsiella pneumoniae* is a growing menace in healthcare settings worldwide (20). This study investigates the comprehensive genomic analysis of a clinical isolate, *Klebsiella* BCSIR-JUMIID, to elucidate its pathogenic and resistance determinants. Whole-genome sequencing and bioinformatic analyses revealed a high-quality assembly with 343 contigs, a total genome length of 5,769,218 bp, and a G+C content of 56.79%, signified by a contig N50 of 32,077 and a contig L50 of 51. These metrics are comparable to other high-quality *K. pneumoniae* genome assemblies, suggesting robust sequencing and assembly protocols (6).

Taxonomic identification, achieved with 100% certainty through whole-genome sequencing and MLST, confirmed the strain as *Klebsiella pneumoniae*. Phylogenetic analysis using the TYGS platform positioned BCSIR-JUMIID within the *K. pneumoniae* family, closely related to strains ATCC 13883 and DSM 30104, indicating a conserved core genome among clinical isolates (7). This phylogenetic proximity underscores the shared evolutionary history and potential for similar pathogenic traits among these strains.

Genome annotation revealed a total of 6,062 coding sequences (CDS), 49 tRNA genes, and 4 rRNA genes, with 1,087 CDS annotated as hypothetical proteins. The high number of functionally assigned proteins (4,975) underscores the bacterium’s extensive metabolic and regulatory capabilities. The presence of multiple proteins with enzyme commission (EC) numbers, gene ontology (GO) assignments, and pathway mappings highlights the complex biochemical landscape of this strain, contributing to its adaptability and survival in diverse environments, including clinical settings (21).

Kaptive analysis identified three main loci: KL150, KL107-D1, and O3b, associated with capsular polysaccharide and O-antigen biosynthesis. The high coverage and identity percentages for KL150 and O3b loci suggest well-preserved regions critical for capsule and O-antigen formation, known virulence factors (8). Conversely, the partial KL107-D1 locus may indicate genetic variability or potential recombination events, reflecting the dynamic nature of these regions.

The genome contained a number of resistance genes encompassing efflux pumps, target modification enzymes, and antibiotic inactivation enzymes, conferring resistance to a broad spectrum of antibiotics (9). The Comprehensive Antibiotic Resistance Database (CARD) analysis highlighted a broad array of antibiotic resistance genes, spanning mechanisms such as efflux pumps (e.g., oqxA and oqxB), beta-lactamases (e.g., SHV-11), and target alteration genes (e.g., parC and gyrA). This aligns with the observed high-level resistance to multiple antibiotic classes, underscoring the significant public health threat posed by this MDR strain (10). The identification of genes like eptB and ArnT, conferring resistance to colistin, is particularly concerning given the critical role of colistin as a last-resort antibiotic (11).

Virulence factor analysis identified genes associated with iron acquisition, adherence, and biofilm formation, suggesting the isolate’s ability to establish successful infections within a host (12). These factors facilitate colonization, biofilm formation, and evasion of host defenses, critical for infection establishment and persistence (13). The presence of these factors underscores the isolate’s pathogenic potential and its capacity to persist in the host environment.

The study also explored horizontal gene transfer (HGT) events, pinpointing mobile genetic elements (MGEs) as potential vehicles for the acquisition of novel resistance genes and virulence factors (14). The detection of numerous MGEs, including plasmids carrying significant resistance genes, underscores the role of HGT in the dissemination of antibiotic resistance (20). The presence of IncFIB(pNDM-Mar) and IncHI1B(pNDM-MAR) plasmids, associated with the NDM gene, highlights the potential for rapid spread of carbapenem resistance (15). This highlights the need for stricter infection control measures to limit the dissemination of resistance determinants within bacterial populations.

Interestingly, the analysis revealed sophisticated defense mechanisms employed by the isolate against bacteriophages (viruses that infect bacteria) (16). The diverse bacteriophage community identified by Phigaro, PHASTER, and VirSorter2 reflects the bacterium’s interactions with its viral predators. The predominance of dsDNA phages indicates their stability and potential role in horizontal gene transfer. The presence of CRISPR-Cas systems, restriction-modification systems, and BREX systems indicates a multi-layered defense strategy against phage predation. Understanding these defense mechanisms might be crucial for the development of novel therapeutic strategies, such as phage therapy, to combat MDR *K. pneumoniae* (17).

The comprehensive genomic analysis of *K. pneumoniae* isolate BCSIR-JUMIID provides valuable insights into its resistance profile, virulence potential, and defense mechanisms. The findings highlight the concerning emergence of MDR strains with a diverse array of virulence factors. Further research is warranted to functionally validate resistance mechanisms, explore phage-bacteria interactions for therapeutic applications, and develop effective strategies to combat this evolving pathogen. This study contributes to the growing body of knowledge regarding the genomic epidemiology of MDR *K. pneumoniae* and underscores the urgent need for integrated One Health approaches to address the complex challenges posed by antimicrobial resistance.

## Conclusion

In conclusion, the genomic characterization of *Klebsiella pneumoniae* BCSIR-JUMIID reveals a strain equipped with extensive resistance mechanisms, a diverse array of virulence factors, and a robust defense system against phage infections. The integration of these findings provides a comprehensive understanding of the pathogenic and antimicrobial properties of this isolate, highlighting its potential public health impact. Future studies are warranted to further explore the functional roles of identified genes and to develop targeted strategies for combating *Klebsiella* infections.

## Acknowledgement

This work was supported by the IAEA Coordinated Research Project (CRP) F23034 (Contract No BGD 23573).

